# Neural signatures of flexible temporal orienting under spatial and motor uncertainty

**DOI:** 10.1101/2025.04.03.646700

**Authors:** Yi Gao, Irene Echeverria-Altuna, Sage E.P Boettcher, Anna C Nobre

## Abstract

The goal-dependent use of temporal expectations enhances visual performance, even without concurrent spatial or motor predictions, yet the underlying neural mechanisms remain unclear. To identify the stages of stimulus processing influenced by temporal orienting of attention, we recorded EEG while participants performed a visual identification task manipulating relevance and temporal predictability of targets. Colored targets appeared in one of two simultaneous visual streams. One color appeared unpredictably while two others appeared predictably early or late. Targets appeared in either stream and required a localization response using the corresponding hand. On each trial, one target was cued as task-relevant. Behaviorally, participants used temporal expectations flexibly to optimize processing of goal-relevant targets. The target-defining cue elicited a contingent negative variation (CNV), modulated by the temporal predictability of anticipated early vs. late targets. Target-related event related potentials (ERP) showed strong modulation by target relevance at multiple stages. Modulations impacted target-selection (N2pc), motor preparation (LRP), and late decision-related factors (P300). Interestingly, only the P300 was additionally modulated by the temporal predictability of targets. The findings reveal how temporal attention can impact multiple stages of stimulus processing through the relevance and temporal predictability of targets even without spatial or motor certainty.

## Introduction

To fine tune perception and behavior within a dynamic and changing environment, the brain prioritizes behaviorally relevant information at key moments in time, a process known as selective temporal attention^1–4^. Orienting of temporal attention can be guided by both the relevance^5–8^ and predictability of event timings^9–14^.

The relevance and prediction of events, linked to goals and expectations, respectively, are separate theoretical constructs^15^, which may impact neural processing in different ways^16,17^. However, in natural settings, they often co-occur and work symbiotically. We consider both aspects, goals and expectations, integral to selective attention – the set of functions that anticipate, select, prioritize, and prepare stimulus contents to guide adaptive behavior^18^. To understand how the brain flexibly uses temporal predictions to focus on goal-relevant signals, we recently designed a new experimental task that separately manipulates goals and temporal expectations derived from learned temporal structures^19,20^.

Visual temporal attention has been proposed to work synergistically with other stimulus attributes, notably spatial locations^21–23^ and motor attributes^24–29^ to facilitate perception and behavior. In light of theoretical proposals for a strong relationship between the neural processing of timing and spatial^30–32^ or motor^33–36^ processing, a reasonable possibility is that temporal orienting of attention does not merely interact with spatial and motor signals but depends upon them. Yet, growing evidence suggests that temporal attention can act without concomitant spatial^37–40^ or action-related^41–43^ expectations.

Of note, Echeverria-Altuna, Nobre and Boettcher^19^ revealed strong behavioral benefits of the flexible allocation of temporal attention in the absence of both spatial and motor certainty. By manipulating both the goal relevance and temporal predictability of targets — two fundamental contributors to attention — the study examined their impact on visual performance. In each trial, three colored targets appeared within rapidly presented visual streams among non-target and masking stimuli. A cue at trial onset designated one color as relevant for responding. Target colors were associated with different temporal predictability. One target color appeared unpredictably, another appeared consistently early, and the other appeared consistently late in the stream. Targets could appear in either of two streams, left or right of visual fixation, and participants responded by indicating the location of the trial-relevant target with the corresponding hand. Despite uncertainty in both the spatial location of targets and the response hand required, target identification was significantly better for temporally predictable targets. The behavioral nature of the task, however, precluded testing the specific stages of neural processing underlying these effects.

The current study, therefore, adapted the experimental task and used electroencephalography (EEG) to test whether goal-dependent attention engages flexible and proactive temporal expectations to anticipate relevant stimuli and to reveal what levels of target processing are modulated. Event-related potentials (ERPs) were derived to assess modulation at different levels of stimulus processing – temporal anticipation, target selection, motor preparation, and decision-related stages.

The electrophysiological signatures of interest were the contingent negative variation (CNV) related to temporal anticipation and motor preparation before a target appears^12,44^; the N2pc, a spatially selective marker of stimulus selection^45–47^; the lateralized readiness potential (LRP), reflecting response preparation or activation^48–52^; and the P300, a late marker of decision formation from sensory information^53,54^.

We hypothesized that behavioral performance would be enhanced for temporally expected compared to temporally unexpected targets, consistent with prior findings^19^. Moreover, we predicted that goal-based signals would interact with temporal predictability to shape proactive anticipation, reflected in a steeper CNV for predictable early versus predictable late targets. A key open question was whether markers of goal-based target selection (N2pc) and later stages of motor preparation and decision-making (LRP, P300) would be influenced by the temporal predictability of targets.

## Methods Participants

The Yale University Institutional Review Board approved the study protocol (2000035997).

Twenty-seven healthy human volunteers participated in the study. Data from two participants were excluded because of poor EEG data quality before any further analysis. The remaining participants had a mean age of 20.64 years (range 18-29 years; 14 female, 11 male). All had normal or corrected vision. Participants provided written informed consent before participation and received monetary compensation for their participation at a rate of $30/h.

The sample size of 25 was set a priori using a simulation-based power analysis^55^ of pilot behavioral data collected using the same task (power = .95).

### Stimuli and task

Participants performed a visual identification task. In each trial, participants saw a color cue followed by two simultaneous rapid-serial-visual-presentation streams, one on the left and one on the right of central fixation. Colored target circles appeared embedded within either stream, and participants had to identify and report the location of the target matching the cue color.

The streams consisted of rapid serial visual presentation of solid-colored distractor and target circles followed by multi-colored masks (**Figure 1a**; see also Echeverria-Altuna et al., 2024). Four colors were used for the distractors and targets (green: #C3CF00, blue: #00ECFF, red: #FF8678, purple: #CAB0FF). Colored circles appeared either on the left or right, accompanied by a white circle (#FFFFFF) of the same dimensions on the other side. Colored and white circles subtended 3.28 degrees of visual angle (DVA) and appeared centered at 5.89 DVA along the horizontal meridian. The purple color was designated as the distractor color for all participants. The other three colors (red, blue, and green) corresponded to possible target colors. Pairs of colored-plus-white stimuli appeared briefly (50 ms) and were immediately masked by multicolored circles appearing at each lateralized location. Masks lasted a variable duration (67-450 ms) until the next pair of solid-colored circles appeared. Thirteen multi-squared, multicolored masks were created by randomly sampling white and the other four colors 20×20 times. The resulting masks were cropped to a circular shape that matched the dimensions of the unicolored circles. One of the thirteen masks was selected randomly for each trial and remained consistent throughout that trial. The duration of both streams was 2250 ms, and participants were instructed to maintain their gaze on the central fixation cross throughout the trial.

**Figure 1.**
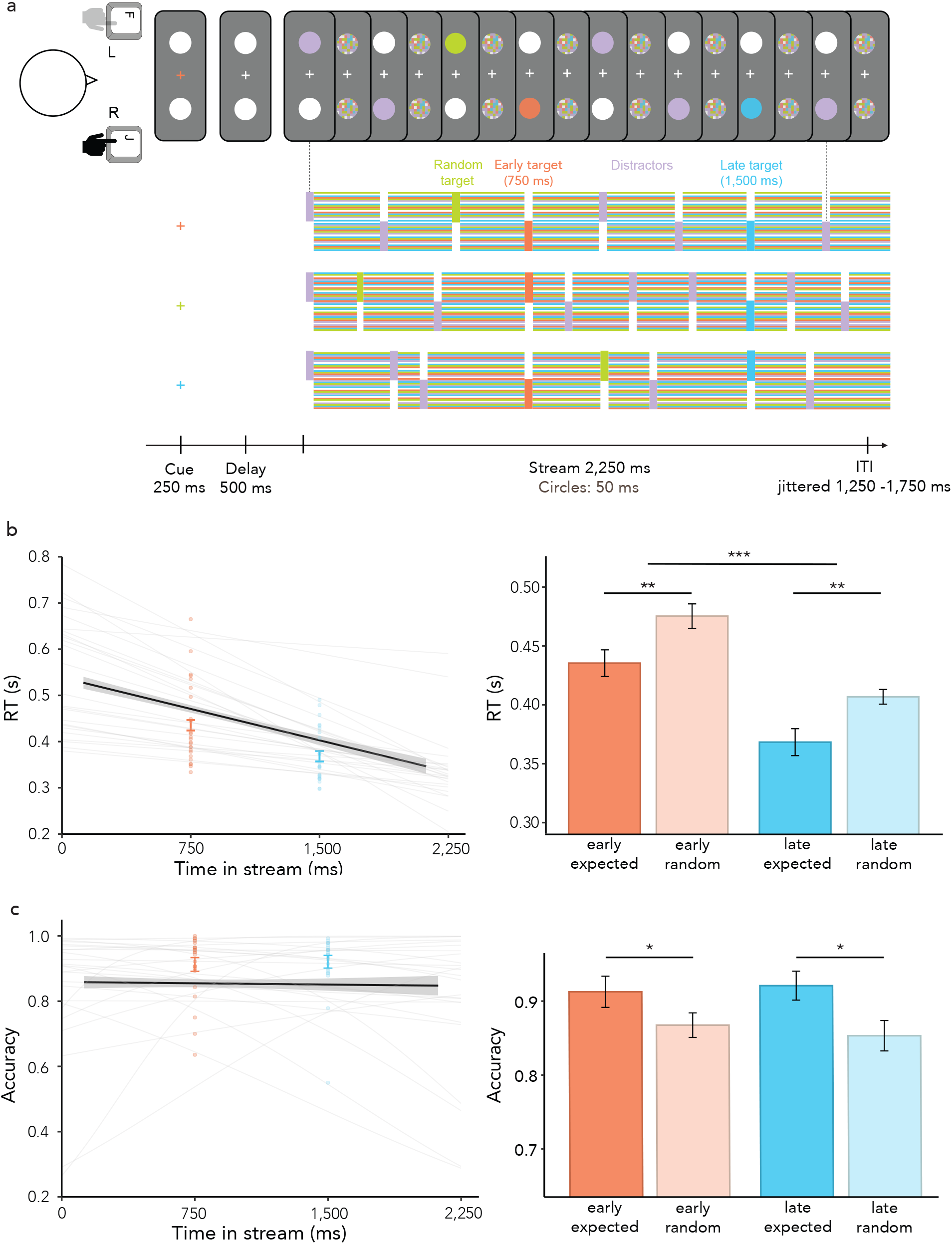
Task design and behavioral results. **(a)** Visual identification task with temporal expectation and task relevance manipulations. A cue at the beginning of each trial — color change of the central fixation cross — indicated the color of the target to detect, out of three possible colors. Participants had to identify the circle with the cued color within one of the two lateralized streams. Upon target identification, participants provided a localization response. Unbeknownst to the participants, the three colored targets differed in their temporal predictability: the early target consistently appeared at 750 ms from stream onset, the late target appeared at 1500 ms from stream onset, and the random target appeared any time throughout the streams (see **Methods**). **(b)** Left: mean RT for random targets as a function of target onset (black line), estimated using participant-specific GLMs (thin grey lines), and mean RTs for early (red) and late (blue) targets across participants. Error bars represent the ±1 standard error of the mean (SEM) and individual points represent data from each participant. Right: mean RTs for early (*M* = 435 ms) and late (*M* = 368 ms) targets across participants, and for random targets at early (*M* = 475 ms) and late (*M* = 407 ms) times (estimated using participant-specific GLMs). **(c)** Left: mean accuracy for random targets as a function of target onset (black line), estimated by participant-specific GLMs (thin grey lines), and mean accuracies for early (red) and late (blue) targets across participants. Error bars represent the ±1 SEM and individual points represent data from each participant. Right: mean accuracies for early (*M* = 91.26%) and late (*M* = 92.09%) targets across participants, and for random targets at early (*M* = 86.75%) and late (*M* = 85.32%) times (estimated using participant-specific GLMs). \**P* < 0.05, \*\**P* < 0.01, \*\*\**P* < 0.001.

Altogether, each stream consisted of nine to twelve colored circles (randomized). To distribute the stimuli evenly across the trials, each of the early (0-750 ms), middle (750-1500 ms), and late (1500-2250 ms) segments of the streams randomly contained three or four colored circles. Purple distractors appeared multiple times throughout the streams. The first colored circle appearing in the streams was always a purple distractor. Each target color appeared up to once during a trial. Trials with all three targets contained six to nine distractors. If one of the three targets was absent, the streams included seven to ten distractors.

At the start of each trial, the fixation cross changed color for 250 ms. This color change served as the cue indicating the target color to be detected in the upcoming visual streams. After a 500-ms delay, the two lateralized streams of white and colored circles began. Colored circles appeared briefly either on the left or right side of the screen, with a white circle displayed on the opposite side. Participants had to identify the target matching the cue color and report its location as quickly as possible. They reported the location of the relevant target by pressing either the leftmost or the rightmost key on the response pad with their left or right hand respectively. Left and right keypresses corresponded to the identification of left- or right-lateralized targets, respectively.

At the end of the 2250-ms streams, the final mask remained on the screen for an additional 1000 ms, followed by trial-specific feedback. Participants received feedback based on their responses: ‘Correct’ for accurate and timely localization responses when the target was present and no responses when the target was absent; ‘Incorrect’ for inaccurate responses or false alarms when the target was absent; ‘Too Early!’ if they responded before the cued target appeared; ‘Too Slow!’ if they responded more than 1000 ms after target onset but before the final mask stopped; or ‘Missing Response!’ if no response was recorded by the end of the final mask.

Each trial was then followed by an inter-trial interval (jittered between 1250-1759 ms), during which the central white fixation cross and two white placeholder circles were displayed on the left and the right side of the screen (5.89 DVA).

The color cue at the beginning of the trial manipulated goal-based attention. The core manipulation of temporal predictability depended on when different target colors were likely to appear during the trial. One color was designated as the early target. When present, it predictably and consistently appeared at a fixed time of 750 ms from the stream onset. Another color was designated as the late target, appearing predictably at a fixed time of 1500 ms from the stream onset. The third color was designated as the random target. It could appear any time from stream onset, except for the time windows around the early (625-875 ms) and late (1375-1625 ms) targets. To minimize foreperiod effects related to mounting temporal conditional probabilities or probability density functions during the trial^56–58^, 50% of random targets appeared in the first segment, 25% in the middle third, and 12.5% in the final third. This meant there was always a 50% chance that a random target would appear in each segment of the trial if it had not yet appeared.

Colors assigned to each target type (early, late, and random) were consistent within a participant and counterbalanced across participants spanning all color-timing condition pairings. All targets had an equal probability of appearing on the left and the right side of the screen, making their appearance spatially unpredictable.

In 62.5% of trials, all three targets (early, late, and random) were displayed. In the remaining 37.5% of trials, two targets were present, and one was absent (12.5% early absent, 12.5% late absent, and 12.5% random absent). In one-third of these target-absent trials (4.17% early absent, 4.17% late absent, and 4.17% random absent), the missing target was also behaviorally relevant (cued). Participants were instructed not to respond in these trials, resulting in a total of 12.5% of trials requiring no response.

Overall, each participant completed 480 trials, 160 in each target-cueing condition – early, late, or random target. Trials from the three conditions were randomly intermixed.

### Procedure

Participants performed the task in person, in an electrically shielded room. They sat approximately 106.5 cm from a 69-cm-wide GIGABYTE M27Q monitor (1,920 × 1,080-pixel resolution; 60-Hz refresh rate). Stimuli were presented using MATLAB 2023a with Psychtoolbox 3.0.19^59^ on a Linux 20.04 operating system. Behavioral data were collected using a LabHackers MilliKey MH-5 USB response box.

Participants completed a practice block of 48 trials before the experimental session and advanced to complete the full task if they reached an average accuracy of 60%. Subsequently, EEG was set up and recorded while participants performed an additional ten blocks of 48 trials. The experimental blocks lasted about 5 minutes each. Participants took short breaks between blocks. In total, the EEG setup took about 40 min, and the experiment session took approximately another 60 minutes to complete.

### Behavioral data analysis

Accuracy measures were restricted to target-present trials. Hits were calculated as the proportion of correct responses when the target was present. The small number of cued target-absent trials per condition (20) precluded sensible analysis. The rates of false alarms in target-absent trials were confirmed to be very low (1.96% of target-absent trials, ranging from 0-5.62%).

For the analyses of response times (RT, time from target onset to correct response), trials were excluded if they had incorrect responses, response times faster than 100 ms, or response times slower than five times the standard deviation of the average RT per participant and condition (early, late, and random). After cleaning, an average of 6.58% (SD: 2.63%) trials were removed per participant from the RT analyses. Accuracy was calculated as the proportion of correct trials out of all trials in each condition.

RTs and accuracy were compared in the three experimental conditions (early, late, and random). To account for potential confounds related to foreperiod effects across conditions, we modeled RT and accuracy for the random target as a linear function of its onset time to establish the baseline level of performance according to stimulus timing alone. The modeling approach was based on a previous related behavioral study in which models with linear and non-linear link functions were compared revealing similar behavioral patterns^19^. For each participant, a generalized linear model (GLM; using the glm function from lme4 R package) was fit to the RTs for random targets assuming a Gamma distribution of RTs and a Gaussian link function^60^. Another GLM was used to model the accuracy of random target responses as a function of their onset assuming a binomial distribution of accuracy and a logit link function. From these participant-specific models, the RT and accuracy values were estimated for the early and late target onset times, respectively. By comparing these estimates with the observed RT and accuracy values when participants responded to early or late targets, we distilled the effects of temporal expectations from foreperiod effects. Finally, 2 × 2 analyses of variance (ANOVAs) were used to assess the effects of predictability (temporally expected vs. random) and time (early vs. late target onsets) on RT and accuracy.

### EEG acquisition and preprocessing

EEG data were acquired using the actiCHamp Plus amplifier and Brain Vision Recorder software (Brain Products GmbH). Electrodes were placed in a 30-electrode montage that followed the international 10-20 system. During the set-up, the impedance of each electrode was lowered to below 5kΩ where possible and to at least 10kΩ in all electrodes. The data were referenced online to the Fz electrode. Additionally, electrodes were placed on the left and right mastoids. Horizontal and vertical eye movements were recorded using two electrodes placed lateral to each eye and two electrodes above and below the left eye, respectively. Data were acquired at a 1000-Hz sampling rate, filtered online between DC and 500 Hz, and saved for offline analysis.

The EEG data were processed and analyzed in Python 3.12.3 and MATLAB 2023a using a combination of custom code, MNE-Python 1.8.0^61^, and SPM12 (Wellcome Department of Imaging Neuroscience, www.fil.ion.ucl.ac.uk/spm). Data were re-referenced to the mean of the left and right mastoids, downsampled to 250 Hz, and band-pass filtered between 0.03 and 50 Hz. Independent Component Analysis (ICA), as implemented in MNE-Python, was applied to identify and remove components related to eye blinks, lateral eye movements, and muscle artifacts. To investigate post-target EEG activity, data were epoched from 200 ms before to 600 ms after target onset, using the 200-ms pre-target window for baseline correction. Epochs with exceptionally high variance were identified and excluded using the generalized extreme studentized deviate (GESD) test^62^. Epochs from trials excluded in the behavioral RT analysis were also removed. On average, 1,256 ± 97 out of 1,440 (87 ± 7%) epochs were retained per participant for further analyses. The continuous, ICA-corrected data were used as the input for a convolution GLM as implemented in SPM12^63^.

### EEG analyses

#### Cue-related CNV

To estimate the CNV leading up to early target appearance in early and late cue trials, the continuous, ICA-corrected data were analyzed using a convolution GLM as implemented in SPM12. The approach minimized interference from overlapping EEG responses to temporally proximal stimuli within the streams. Deconvolved evoked responses were modeled from 500 ms before to 1500 ms after events corresponding to cues (early, late, and random), targets (early, late, and random; left and right; cued and uncued), distractors, and responses (left and right hand-response to early, late, random targets). Only the trials retained from the behavioral RT analysis were modeled. Cue-locked responses were baseline corrected by subtracting the average potential in the 200-ms window preceding cue onset. To isolate the CNV, we focused on the dynamics at a predefined electrode cluster (Fz, FC1, FC2, Cz), consistent with extensive previous work showing that the CNV is more pronounced at frontocentral sites^64^, and a time window between cue onset and the appearance of the early target. Data were averaged across the selected electrodes and compared between conditions in which an early or late target was cued. We focused on the CNV elicited when early vs. late targets were cued to maximize the trial numbers and compare the neural modulation related to clearly distinct temporal expectations.

#### Target-related ERPs

We applied a surface-Laplacian transform to the epoched EEG data, as implemented with MNE-Python, to reduce the effects of volume conduction^65^. Target-locked ERPs were computed for early, late, and random targets in trials when they were cued (task-relevant) and uncued (task-irrelevant). For task-irrelevant targets, only trials in which the cued target had not yet appeared were included, eliminating contamination from previous responses and task disengagement.

To investigate the modulation of the N2pc by target relevance, EEG activity was analyzed at the left (P7, O1) and right (P8, O2) posterior sites where the N2pc is typically observed^66^. PO7/PO8 were unavailable in the montage. The N2pc measurements were obtained by subtracting ipsilateral activity (average of left-sided electrodes for left targets and right-sided electrodes for right targets) from contralateral activity (average of left-sided electrodes for right target and right-sided electrodes for left targets) at the respective electrode sites.

To analyze the LRP, we used the C3 and C4 electrodes positioned over motor cortical areas^67^. Readiness potentials were calculated by comparing potentials at contralateral versus ipsilateral electrodes relative to the response hand, yielding a single LRP waveform per target relevance condition.

The centroparietal electrodes (CP1, CP2, P3, Pz, P4) were used for the P300 analysis, based on previous work of P300 in the context of perceptual decision-making^68^. Data were averaged across the selected electrodes and compared across trials in which targets — early and late — were task-relevant or task-irrelevant.

Topographies of N2pc and LRP were calculated as average contra-vs. ipsilateral contrasts over 210-260 ms and 250-350 ms time windows, respectively, across all sensor pairs. Topographies for P300 represented the evoked data from all sensors averaged across the 350–450 ms post-target window.

Moreover, the N2pc, LRP, and P300 time-courses locked to task-relevant targets were collapsed across the temporally predictable (early and late) conditions and compared to the temporally unpredictable (random) condition. To minimize potential foreperiod effects, random targets were sampled from the first, second, and third segments of the stream at a 1:2:1 ratio, equating the average target onset between conditions.

#### Relationship between EEG and behavior

To investigate the relationship between the observed target-evoked EEG activity and RT, the behavioral measure most strongly modulated by temporal expectations in this task, we employed the following approach. We grouped participants based on their RT benefit of temporal expectation, defined as the average RT difference between expected and random targets at early and late time points (see **behavioral data analysis**). Participants were split into high and low RT benefit groups using the median RT benefit across participants for each trial type: early and random task-relevant target responses were grouped based on the RT benefit at the early time point, while late and random task-relevant target responses were grouped based on the RT benefit at the late time point. We then compared the target-locked potential that was most strongly modulated by temporal predictability (i.e., P300; **Figure 4**) between these groups, using cluster-based permutation testing, as detailed below.

#### Statistical evaluation

We used cluster-based permutation testing to evaluate the baseline-corrected evoked EEG responses between experimental conditions, allowing for robust statistical evaluation along the time axes while circumventing the multiple-comparisons problem^69^. Temporal clusters across the participant-specific evoked time courses were statistically assessed using 1000 permutations and a significance threshold set at an alpha level of .05. To evaluate the CNV between conditions (cue early vs. late), we focused on the period just before the onset of early targets (1200 – 1500 ms; i.e., 300 ms before early target onset). Since the streams always consistently started at 750 ms after cue appearance, this interval homed in on the time of putative differential temporal expectations and minimized the potential contamination from the evoked responses related to the onset of the visual streams.

All statistical analyses were applied to the EEG time courses extracted from the predefined electrode clusters. Topographical analyses were not subjected to further inferential statistical tests but were visualized to confirm the suitability of *a priori* electrode selections and the physiological plausibility of the identified patterns^70^.

## Results

### Behavioral benefits

A 2 × 2 ANOVA of RT revealed a main effect of predictability (expected vs random; *F*(1, 24) = 10.82, *p* = .003, *η*^*2*^_*G*_ = .06) and a main effect of time (early vs late; *F*(1, 24) = 41.24, *p* < .001, *η*^*2*^_*G*_= .16) with no interaction between the factors (*F*(1, 24) = .01, *p* = 0.92, *η*^*2*^_*G*_ = .00); **Figure 1b**). For accuracy, we found a main effect of predictability (*F(*1, 24) = 4.89, *p* = .037, *η*^*2*^_*G*_ = .06), with no main effect of time (*F*(1, 24) = 0.03, *p* = .86, *η*^*2*^_*G*_ = .00) or interaction between the factors (*F*(1,24) = .72, *p* = .40, *η*^*2*^_*G*_ = .00; **Figure 1c**). Therefore, participants were significantly faster and more accurate at detecting temporally expected targets compared to temporally unexpected targets.

### Cue-related CNV modulation

Focusing on the interval between cue and target onset, we observed a gradual emergence of a negative potential at the medial frontal scalp (i.e., CNV) in both cue-early and cue-late conditions. The potential was slightly more negative when the early target was cued versus the late target. This difference between conditions was significant during the 300-ms window immediately before early target onset (**Figure 2a** black line; cluster *p* = .037, one-tailed). The topography of the difference between conditions confirmed the frontocentral distribution (**Figure 2b**).

**Figure 2.**
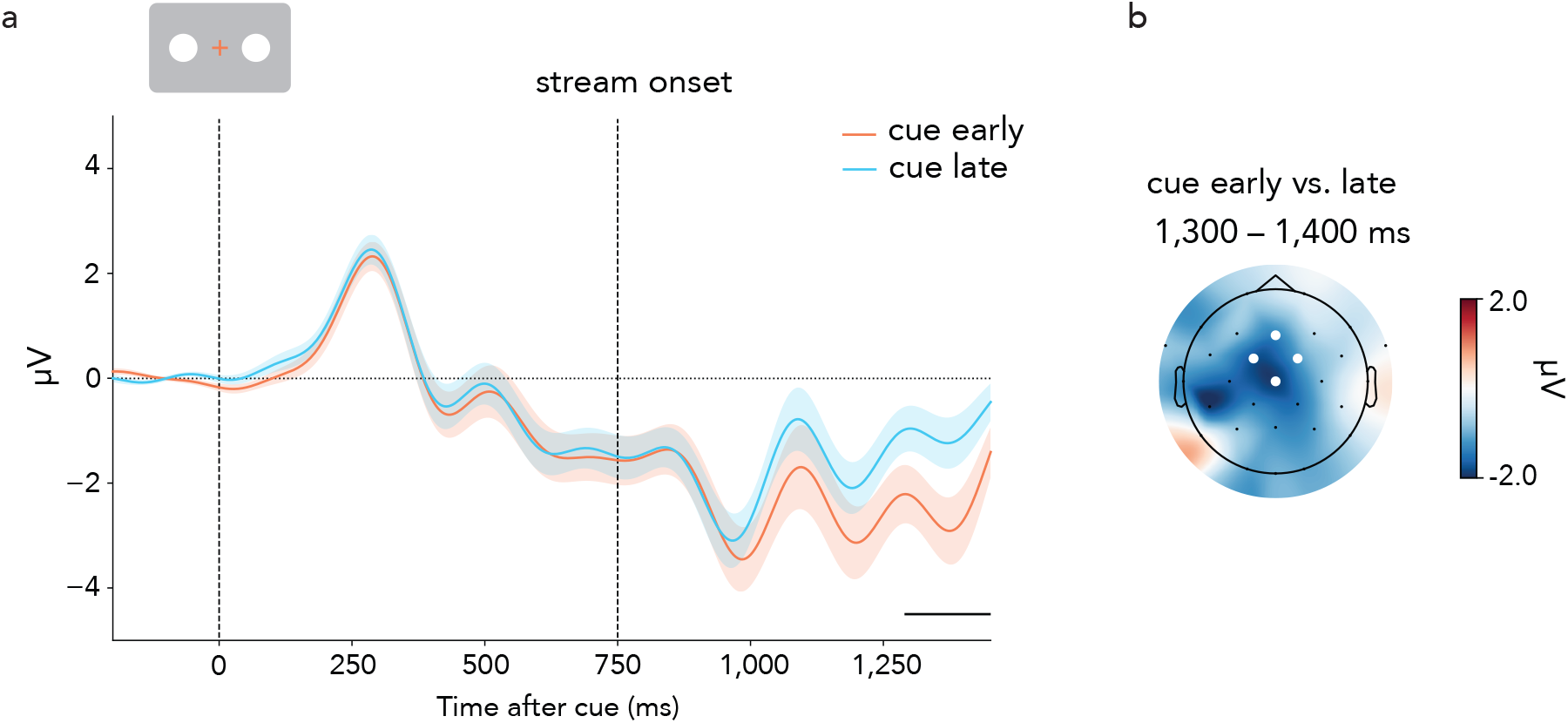
CNV associated with temporal expectations during the pre-target period. **(a)** Average EEG potentials in the selected frontal-central electrodes (marked in B; [Fz, FC1, FC2, Cz]) locked to cue onset. The CNV is observed as a gradual increase in the signal negativity between cue and target onset. Dashed vertical lines represent, in sequential order, the cue onset and the onset of the streams. The black line below the time courses indicates a significant difference between conditions (early vs. late) after a cluster-based permutation analysis in the 1200-1500 ms post-cue window, immediately before early target onset. Shaded areas indicate ± 1 SEM calculated across participants (*n* = 25). **(b)** The topography associated with the difference between conditions shown in A, averaged over 1300-1400 ms after cue.

### Modulation of target-related ERPs

#### Modulation of N2pc

Analysis of the N2pc tested the effects of goal relevance on target selection based on its defining sensory attributes. **Figures 3a-c** show the lateralized ERP activity in the predefined posterior electrodes, relative to the location of the target. The N2pc was clearly identified as a stronger negative deflection at electrodes contralateral compared to ipsilateral to the target location between ~ 200 and 280 ms. Across early, late, and random targets, a clear difference was observed in the contralateral-minus-ipsilateral trace for cued, task-relevant targets compared to uncued, task-irrelevant targets. The N2pc elicited by the early target was significantly larger when the early target was relevant vs. irrelevant in either late (**Figure 3a**, black line; cluster *p*_*cue early vs. late*_ = .005) or random (**Figure 3a**, grey line; cluster *p*_*cue early vs. random*_ = .002) trials. Similarly, the N2pc elicited by the late target was significantly larger when the late target was relevant vs. irrelevant in random trials (**Figure 3b**, grey line; cluster *p*_*cue late vs. random*_ = .002). In random trials, the N2pc was also significantly larger when random targets were relevant vs. irrelevant in either early (**Figure 3c**, black line; cluster *p*_*cue random vs. early*_ = .006) or late (**Figure 3c**, grey line; cluster *p*_*cue random vs. late*_ = .005) trials. For early and random targets, the evoked potentials did not differ between the two irrelevant conditions. As explained in the Methods, only trials for which the target had not yet appeared were included in the irrelevant target conditions. For this reason, it was not possible to analyze late target responses in early trials. Topographical plots confirmed the lateralized posterior distribution of the N2pc effect. In the late task-relevant vs. irrelevant topography, we also observed lateralized neural activity in central sites, which may reflect the preparation of motor responses towards the end of the trial.

**Figure 3.**
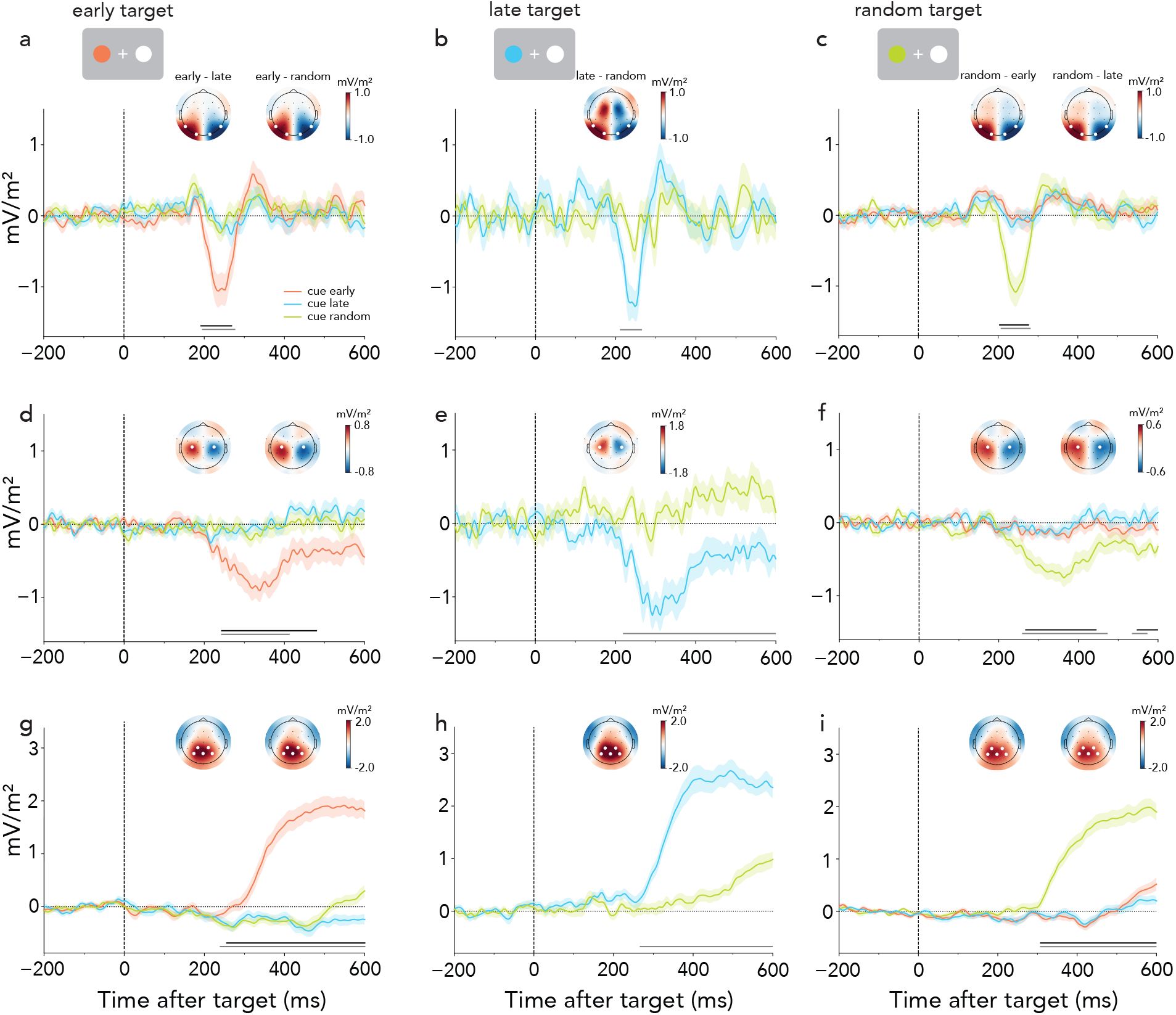
Task-relevant, temporally expected targets elicit N2pc, LRP, and P300 signals. **(a, b, c)** N2pc waveforms (contralateral minus ipsilateral ERPs relative to the side of the relevant target) in selected parietal-occipital electrodes ([P7, P8, O1, O2], marked in the topographies) as a function of task relevance (cued vs. uncued) for early **(a)**, late **(b)**, and random **(c)** targets, along with the associated topographies averaged across 210-260 ms. **(d, e, f)** LRP waveforms (contralateral minus ipsilateral ERPs relative to the response hand) recorded from selected central electrodes ([C3, C4], marked in the topographies) are shown as a function of task relevance (cued vs. uncued) for early **(d)**, late **(e)**, and random **(f)** targets, along with the associated topographies averaged across 250-350 ms. **(g, h, i)** P300 waveforms in selected central-posterior electrodes ([CP1, CP2, P3, Pz, P4], marked in the topographies) as a function of task relevance (cued vs. uncued) for early **(g)**, late **(h)**, and random **(i)** targets, along with the associated topographies averaged across 350-450 ms. Dashed lines represent target onset. Horizontal lines indicate statistically significant clusters (black for the difference between cue-early and cue-late conditions, grey for the difference between cue-early/late and cue-random conditions). Shaded areas in all plots indicate ± 1 SEM calculated across participants (*n* = 25).

#### Modulation of LRP

Analysis of the LRP tested for effects of goal relevance on motor preparation. **Figures 3d-f** show the differences in ERP activity contralateral vs. ipsilateral to the side of the responding hand, extracted over the selected central electrodes, separately for early, late, and random targets.

A direct comparison between relevant and irrelevant target conditions revealed significant differences in the LRP after the onset of relevant vs. irrelevant targets. The LRP elicited by the early target was significantly larger when the early target was relevant than when it was irrelevant in either late (**Figure 3d**, black line; cluster *p*_*cue early vs. late*_ = .003) or random (**Figure 3d**, grey line; cluster *p*_*cue early vs. random*_ = .007) trials. Similarly, the LRP elicited by the late target was significantly larger when the late target was relevant than when it was irrelevant in random trials (**Figure 3e**, grey line; cluster *p*_*cue late vs. random*_ = .001). In random trials, the LRP was also significantly enhanced when random targets were relevant vs. irrelevant in either early (**Figure 3f**, black lines; clusters *p*_*cue random vs. early*_ = .002, .036) or late (**Figure 3f**, grey lines; clusters *p*_*cue random vs. late*_ = .001, .048) trials. For early and random targets, the evoked potentials did not differ between the two irrelevant conditions. Only trials for which the target had not yet appeared were included in the irrelevant target conditions. Topographical plots confirmed the lateralized central distribution of the LRP effects, consistent with modulation of motor activity.

#### Modulation of P300

Analysis of the P300-like late positive potential tested for effects of goal relevance on late, decision-related factors. Across early, late, and random targets, late central-posterior ERP activity was significantly enhanced for task-relevant compared to task-irrelevant targets (**Figures 3g-i**). The P300-like potential elicited by the early target was significantly larger when the early target was relevant than when it was irrelevant in either late (**Figure 3g**, black line; cluster *p*_*cue early vs. late*_= .001) or random (**Figure 3g**, grey line; cluster *p*_*cue early vs. random*_ = .001) trials. Similarly, the P300-like potential elicited by the late target was significantly larger when the late target was relevant than when it was irrelevant in the random condition (**Figure 3h**, grey line; cluster *p*_*cue late vs. random*_= .001). In random trials, the P300 was also significantly larger when the random targets were relevant vs. irrelevant in either early (**Figure 3i**, black line; cluster *p*_*cue random vs. early*_ = .001) or late (**Figure 3i**, grey line; cluster *p*_*cue random vs. late*_ = .001) trials. Again, for early and random targets, the potentials did not differ between the two irrelevant conditions until after 500 ms. Only trials for which the target had not yet appeared were included in the irrelevant target conditions. The significant clusters indicating P300 modulation began around 300 ms, overlapping with the period of the LRP modulation. Topographical plots confirmed the central-posterior distribution of the effects.

#### Effects of temporal expectation on target-related ERPs

To assess which neural markers of goal-based selection were influenced by temporal expectation, we compared lateralized ERP components (N2pc and LRP) and P300 responses between task-relevant random (evenly sampled across the trial) and expected targets (early and late trials combined). No significant differences emerged for N2pc or LRP (**Figures 4a-b)**. In contrast, the P300-like potential was significantly larger for temporally expected targets compared to random targets (**Figure 4c**, black line; cluster *p* = .006).

**Figure 4.**
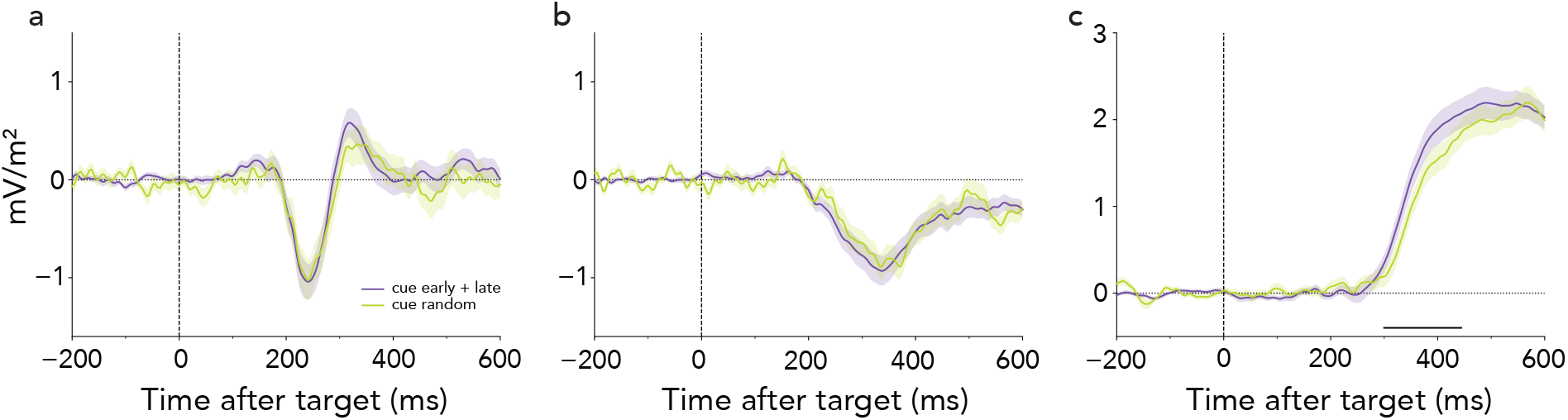
**(a)** N2pc, **(b)** LRP, and **(c)** P300 waveforms as a function of target predictiveness, comparing temporally predictable (early and late combined) and unpredictable (random) targets. Random target trials are sampled at a 1:2:1 ratio across the first, second, and third segments of the streams. Dashed lines represent target onset. Horizontal lines indicate statistically significant clusters between cue-expected (early and late) and cue-random conditions. Shaded areas in all plots indicate ± 1 SEM calculated across participants (*n* = 25).

#### Target-related ERP modulations relate to behavioral benefits of temporal attention

We next investigated whether the effects of the temporal predictability of targets on P300 were linked to the behavioral benefits of temporal attention (see **Figure 1**). Specifically, we compared the P300 between participants with high vs. low behavioral benefits of temporal expectation (see **Methods**). As shown in **Figure 5**, participants with a stronger behavioral benefit displayed a larger P300 for early targets (cluster *p*_*high vs. low benefit at early time*_ = .014; **Figure 5a**) but not random targets (**Figure 5b**). These findings suggest that the late stage of target processing is closely linked to the behavioral benefits of visual temporal expectation.

**Figure 5.**
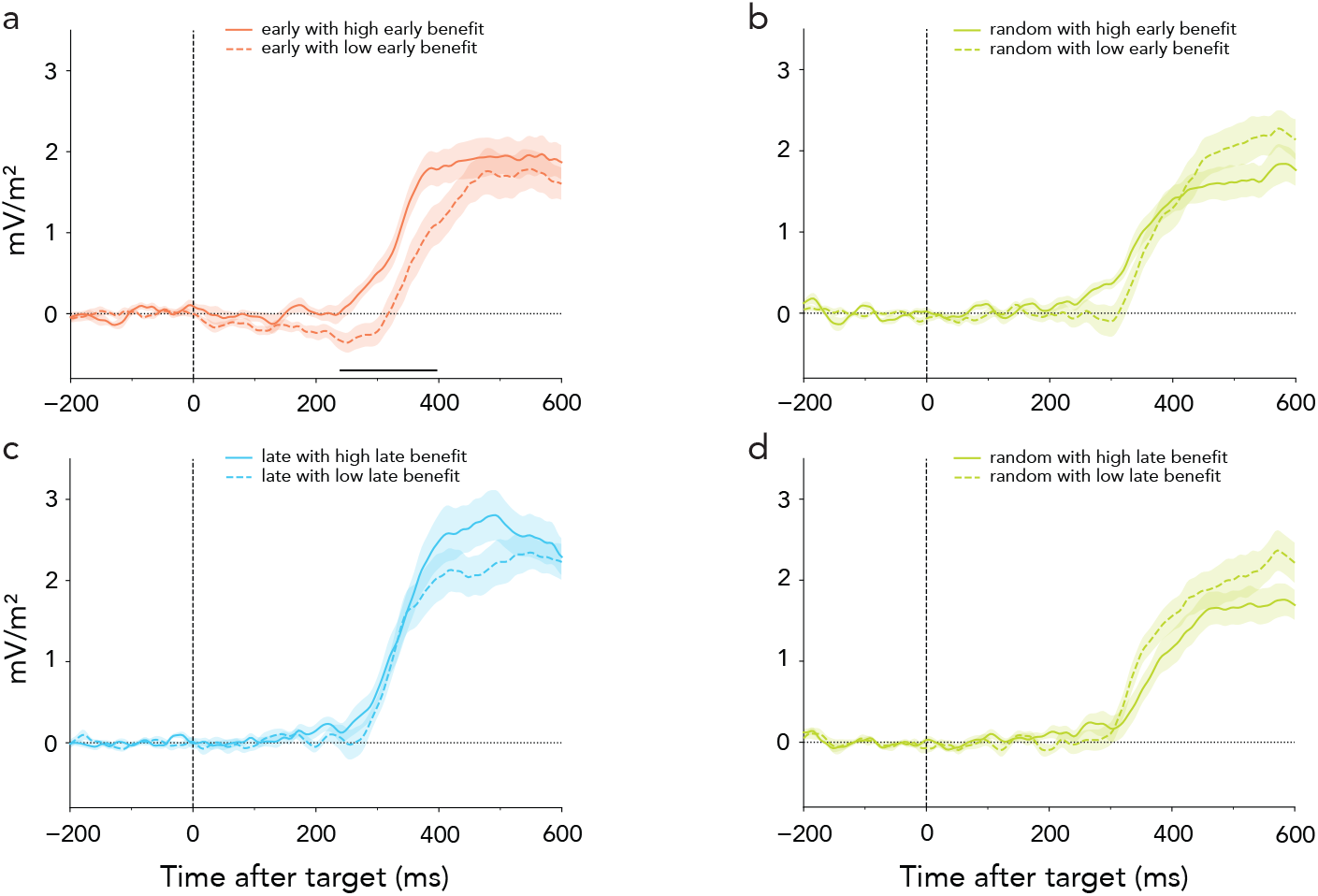
P300 activity is linked with the behavioral benefit of temporal expectations. P300 waveforms are shown for target-relevant trials with high (solid lines) and low (dashed lines) behavioral benefit of temporal expectations, based on a median split of RT differences between expected and random targets at the early **(a, b)** and late **(c, d)** time points, computed per participant (see **Methods**). Significant clusters from cluster-based permutation tests are indicated with horizontal black lines. The vertical dashed lines represent target onset. Shaded areas in all plots indicate ± 1 SEM calculated across participants (*n* = 13 for high benefit group, *n* = 12 for low benefit group).

Although P300 amplitudes for late targets did not systematically differ between high- and low-benefit trials, a trend toward stronger modulation in the high-benefit group was observed (**Figure 5c**). In contrast, if anything, P300 responses to random targets showed greater modulations in the low-benefit group (**Figure 5d**).

## Discussion

The behavioral and neural results show that goal-based attention capitalizes on incidentally learned temporal predictions to guide behavior.

The behavioral benefits of detecting temporally predicted vs. unpredicted targets more accurately and more quickly replicated previous findings^19^. As in the previous study, performance also revealed the contributions from additional effects related to the passage of time in the trial (foreperiod). In the current study, the foreperiod effect was restricted to reaction times, which improved over the course of the trial. Accuracy, in contrast, was consistent over time. The foreperiod effects on response times are intriguing given that the conditional probability of random targets was similar across early, middle, and late segments of the trial. They suggest that the probability density function may provide a better predictor of temporal expectation benefits^57^ and/or that effects related to increasing general preparation also contribute to performance.

Neural markers of temporal anticipation were modulated by the flexible use of incidentally learned temporal predictions. The central cueing stimulus elicited a characteristic CNV potential, previously associated with target anticipation and temporal expectation^8,12^,71. The amplitude of the CNV was modulated by temporal expectation for the target. Following the onset of the stimulus streams, the potential became more negative when the early-predicted target had been cued compared to the late-predicted target. Notably, the effect started after the temporally predictable onset of the two stimulus streams, when the differential temporal predictions of the different colored targets became relevant.

Target-locked EEG markers showed strong goal-driven modulation of successive processing stages – target selection, associated motor preparation, and late evaluative processes. Task-relevant, cued targets elicited strong, lateralized N2pc potentials over the posterior scalp, indicating spatial selection of the defining target feature^45,47^. In contrast, non-relevant targets elicited little or no lateralized activity during the same period, and did not differ from one another. This pattern of strong N2pc for the relevant target and equivalent absent potentials for the irrelevant targets was observed across early, late, and random targets. There was no clear modulation of the goal-based target selection process by temporal expectation in the task. Putative benefits of temporal expectation were difficult to assess because predictable and unpredictable targets occurred at different time points in the trial, so conclusions must remain tentative. However, no differences were observed when comparing the N2pc potentials elicited by predictable relevant targets at early and late time points with random relevant targets drawn across early, middle, and late segments of the stream. Additionally, there was no evidence for selective suppression of task-irrelevant targets by temporal expectations: early target-evoked potentials were equivalent whether participants expected late targets (focused temporal expectations), or random targets (more distributed temporal expectations).

Following the N2pc, strong lateralized central LRPs emerged, showing a similar pattern: task-relevant, cued targets elicited robust LRPs, whereas the same stimuli produced no LRPs when task-irrelevant. No differences were noted for task-irrelevant early target stimuli, regardless of whether participants were cued to detect a predictable (late) or unpredictable (random) target. Surprisingly, comparing the LRPs elicited by temporally predicted and unpredicted targets showed no significant differences.

Goal-based attention strongly modulated the late positive P300-like potential associated with late-stage processes like stimulus evaluation and decision-making^53,54^. Again, the potential was exclusively elicited by task-relevant targets, with no differences between different task-irrelevant conditions. Comparing temporally predictable and unpredictable targets yielded evidence that temporal expectation influenced this stage of processing: the P300 was larger for temporally anticipated targets. The findings echoed previous reports of P300 modulation by temporal expectation, in which the P300 starts earlier and has larger amplitude for temporally anticipated relevant events^8,12^,72. The P300 potential scaled with individual behavioral benefits of temporal expectation – larger P300 responses were observed for valid targets in participants with greater RT gains, while responses to random targets were unaffected. These results further suggest that behavioral benefits of temporal expectation arise from prioritized processing of expected targets, rather than diminishing processing of unpredictable ones. Future studies including invalid targets may help further disentangle the directionality of these effects. The lack of modulation at the LRP stage suggests that temporal expectation influences a late process following response selection or preparation. However, the rise time of the P300 relative to the response time leaves open a possibility that other aspects of the response process is facilitated. Alternatively, decision-related processes concurrent with or following response-related processes may be enhanced^53,73^.

The limited influence of temporal expectation on early markers of goal-based attention in our task was surprising. Speculatively, the absence of a modulation of lateralized markers of target processing (N2pc) and motor preparation (LRP) may be related to the purposeful manipulation of spatial and motor uncertainty in the present study. In the current design, we employed complex visual streams with temporally jittered distractors and masks, allowing us to probe how temporal expectations operate in dynamic and attentionally demanding environments. This complexity highlights the adaptability of the attentional system but may have limited our ability to detect early sensory components leading up to target selection—such as the P1, N1, and the selection negativity associated with color-based attention^74,75^. Future work could revisit the influence of temporal expectation on early perceptual processing using task designs with reduced visual interference. In the absence of earlier markers of attention modulation or color-based selection in the task, our results add support to the notion that spatial and motor attributes serve as key channels for carrying or potentiating the effects of temporal expectation^3,72^.

Notably, our design minimized spatial and motor competition by presenting targets in opposite hemifields and assigning responses to two different hands. Whether temporal predictions confer strong performance benefits under conditions of much stronger spatial or motor competition remains an open question. Finally, while the present study showed that temporal expectations facilitated goal-based target selection according to color, it will be interesting to test whether the effects could extend to other stimulus features like orientation^76^, shape, or motion.

Overall, our findings extend previous observations that temporal predictions can enhance goal-based attention performance even in the absence of spatial or motor certainty^19^. They reveal the engagement of flexible, proactive temporal anticipation effects but suggest that modulatory effects on target processing may occur at late stages. Future work should revisit the possibility of early-stage target-related modulation and further explore the boundary conditions for observing temporal expectation without spatial or motor certainty.

## Author Contributions

Yi Gao: Conceptualization, Methodology, Software, Investigation, Formal analysis, Visualization, Writing – original draft. Irene Echeverria-Altuna: Conceptualization, Methodology, Writing – review & editing. Sage E.P. Boettcher: Conceptualization, Methodology, Writing – review & editing. Anna C. Nobre: Conceptualization, Methodology, Writing – review & editing, Funding acquisition, Supervision.

## Data availability

Data and analysis scripts are available from the corresponding author, Yi Gao, upon reasonable request.

